# Proinflammatory responses in SARS-CoV-2 infected and soluble spike glycoprotein S1 subunit activated human macrophages

**DOI:** 10.1101/2021.06.14.448426

**Authors:** Kim Chiok, Kevin Hutchison, Lindsay Grace Miller, Santanu Bose, Tanya A. Miura

## Abstract

Critically ill COVID-19 patients infected with SARS-CoV-2 display signs of generalized hyperinflammation. Macrophages trigger inflammation to eliminate pathogens and repair tissue, but this process can also lead to hyperinflammation and resulting exaggerated disease. The role of macrophages in dysregulated inflammation during SARS-CoV-2 infection is poorly understood. We used SARS-CoV-2 infected and glycosylated soluble SARS-CoV-2 Spike S1 subunit (S1) treated THP-1 human-derived macrophage-like cell line to clarify the role of macrophages in pro-inflammatory responses. Soluble S1 upregulated TNF-α and CXCL10 mRNAs, and induced secretion of TNF-α from THP-1 macrophages. While THP-1 macrophages did not support productive SARS-CoV-2 replication, virus infection resulted in upregulation of both TNF-α and CXCL10 genes. Our study shows that S1 is a key viral component inducing inflammatory response in macrophages, independently of virus replication. Thus, virus-infected or soluble S1-activated macrophages may become sources of pro-inflammatory mediators contributing to hyperinflammation in COVID-19 patients.

## 1. Introduction

Severe Acute Respiratory Syndrome Coronavirus 2 (SARS-CoV-2) is the causative agent of Coronavirus Disease 2019 (COVID-19). Severely ill COVID-19 patients display lung tissue damage associated with cell death and pathologic inflammation^1,2^ linked to enhanced pro-inflammatory cytokine and chemokine levels (*e.g.*, TNF-α and CXCL10)^3,4^. These pathologies are compatible with dysregulated inflammatory response characteristic of cytokine release syndrome or macrophage-activation syndrome^5^ and generalized hyperinflammation^6^. These patients often progress to respiratory failure due to complications from hyperinflammation and require mechanical ventilation. Analysis of bronchoalveolar lavage fluid (BALF) from critically ill COVID-19 patients revealed upregulation of inflammatory cytokine signatures, indicating influx of active inflammatory macrophages in the airways^7,8^. Single-cell RNA sequencing also detected SARS-CoV-2 RNA in BALF-associated macrophages^8^, whereas virus antigen was detected in subsets of tissue-resident and lymph node-associated macrophages^9^. Macrophages mediate inflammatory responses following infection, migrating to affected tissue to eliminate invading pathogens. These findings implicate macrophages in the exaggerated inflammatory response during SARS-CoV-2 infection. In this study, we analyzed pro-inflammatory response against SARS-CoV-2 and soluble SARS-CoV-2 spike glycoprotein S1 subunit in the human-derived THP-1 macrophage cell-line.

## 2. Results and Discussion

### 2.1. Soluble SARS-CoV-2 spike protein S1 subunit induces inflammatory response in THP-1 cells

Reports using non-soluble SARS-CoV-2 trimeric spike (S) glycoprotein^10^ and purified S1 subunit produced in *E.coli*^11^ suggest that the S protein activates inflammation in macrophages. Prefusion trimeric S is cleaved by cellular proteases^12^ that dissociate the S1 subunit during virion assembly^13^ or after engagement of the ACE2 receptor^14^. Trimeric S is transient on the surface of virions, and constructs designed to stabilize this conformation do not reflect the dynamic state of the S glycoprotein and subunits in contact with cells. Importantly, dissociated S1 may remain engaged to cell receptors and stimulate yet undefined effects. Furthermore, S1 is glycosylated at numerous positions^15^ that mediate functions such as shielding of viral epitopes. Glycosylation patterns are not recapitulated in *E.coli* used to purify proteins and thus, non-glycosylated S1 produced in *E.coli* may not reproduce the biological effects of SARS-CoV-2 S1.

To clarify the role of S1 in proinflammatory responses in macrophages, we tested whether glycosylated, soluble SARS-CoV-2 S1 produced in mammalian cells induced expression of pro-inflammatory and antiviral cytokines. We evaluated expression of the pro-inflammatory cytokines TNF-α, CXCL10 and IFN-γ^16,17^ due to their association with hyperinflammation in COVID-19 patients, and the antiviral cytokine IFN-β since it restricts SARS-CoV-2 infection^16^. While S1 did not induce gene expression of antiviral IFN-β (Fig.1A) or IFN-γ (Fig.1B) in THP-1 macrophages, expression of proinflammatory TNF-α (Fig.1C) and CXCL10 (Fig. 1D) was upregulated following exposure to S1. TNF-α was upregulated by 30-fold at 4h post treatment with S1 (*p* < 0.05) and remained upregulated by two-fold at 16h post treatment (Fig. 1C). CXCL10 expression was consistently upregulated by 3 to 8-fold in THP-1 macrophages exposed to S1 up to 16h post treatment (*p*<0.05) (Fig. 1D). Although macrophages respond to IFN-γ by producing CXCL10, they are not substantial sources of IFN-γ^18^, which is mostly produced by lymphocytes to recruit macrophages to infection sites^19^. Thus, SARS-CoV-2 S1 upregulated CXCL10 independently of IFN-γ, similar to LPS^20^ and TNF-α^21^.

**Figure 1.**
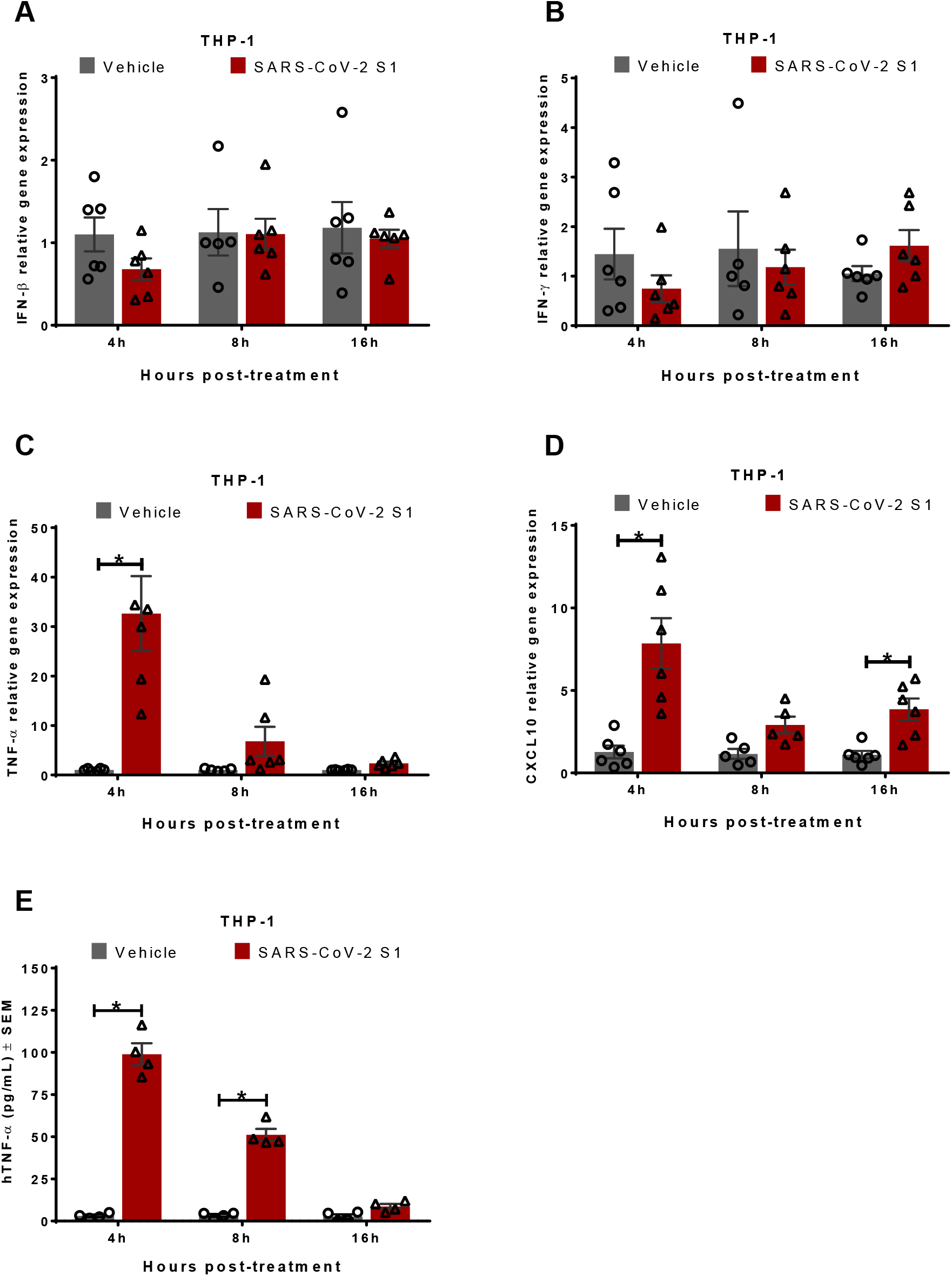
SARS-CoV-2 soluble Spike protein S1 subunit (S1) induces inflammatory response in THP-1 macrophages. THP-1 cells were treated with purified recombinant soluble S1 protein (0.6ug/mL, 8nM) or vehicle for indicated times. RT-qPCR was used to detect relative gene expression of cytokines IFN-β (A), IFN-γ (B), TNF-α (C), and CXCL10 (D). (E) Secretion of TNF-α was determined by ELISA assays in supernatants from THP-1 cells treated with purified recombinant soluble S1 protein. Error bars denote the standard error of the mean (SEM) from 3 biologically independent experiments. \**p*< 0.05, \*\**p*<0.01 determined by Two-way ANOVA adjusted by Sidak’s multiple comparison test.

We further examined release of TNF-α from THP-1 macrophages treated with the SARS-CoV-2 S1 subunit. Treatment with S1 increased secretion of TNF-α in macrophages up to 30 times more than vehicle-treated controls at 4h post treatment (*p*< 0.05) (Fig. 1E). These results demonstrated that soluble S1 alone suffices to activate inflammatory response in macrophages independently of full-length S protein, S-trimers and virus infection. Non-glycosylated S1 from *E.coli* induced TNF-α secretion in murine macrophages^11^. Although these studies were conducted in mouse macrophages^11^, non-glycosylated S1 may expose sites able to trigger pro-inflammatory response, which are cryptic in the glycosylated S1 protein. By utilizing physiologically relevant glycosylated S1 derived from mammalian cells, we demonstrate its pro-inflammatory activity in human macrophages.

### 2.2. Inflammatory response in SARS-CoV-2 infected THP-1 macrophages in the absence of productive viral replication

Since SARS-CoV-2 S1 induced pro-inflammatory response in macrophages independently of virus infection or replication (Fig. 1), we next evaluated pro-inflammatory response and viral replication in THP-1 macrophages infected with SARS-CoV-2. As expected, infection with SARS-CoV-2 in susceptible Vero E6 cells led to exponential increase of viral nucleocapsid (N) (Fig. 2A) and S (Fig. 2B) genes. In contrast, expression of N (Fig. 2C) and S (Fig. 2D) genes was reduced in THP-1 macrophages over the course of infection. Virus replication and release of infectious progeny was determined by TCID_50_ assay in supernatants from SARS-CoV-2 infected cells to corroborate viral RNA findings. While infected Vero E6 cells supported robust release of infectious virions due to productive replication, infectious virus production from infected THP-1 cells became undetectable after 8h post-infection (Fig. 2E). Finally, THP-1 macrophages did not develop cytopathic effects (CPE) following SARS-CoV-2 infection, whereas Vero E6 cells displayed progressive monolayer damage over the course of infection (Fig. 2F). These results indicate that THP-1 human macrophages do not support productive replication of SARS-CoV-2. Our study is in accord with a report showing non-productive replication of SARS-CoV-2 in human monocyte-derived macrophages and DCs^22^.

**Figure 2.**
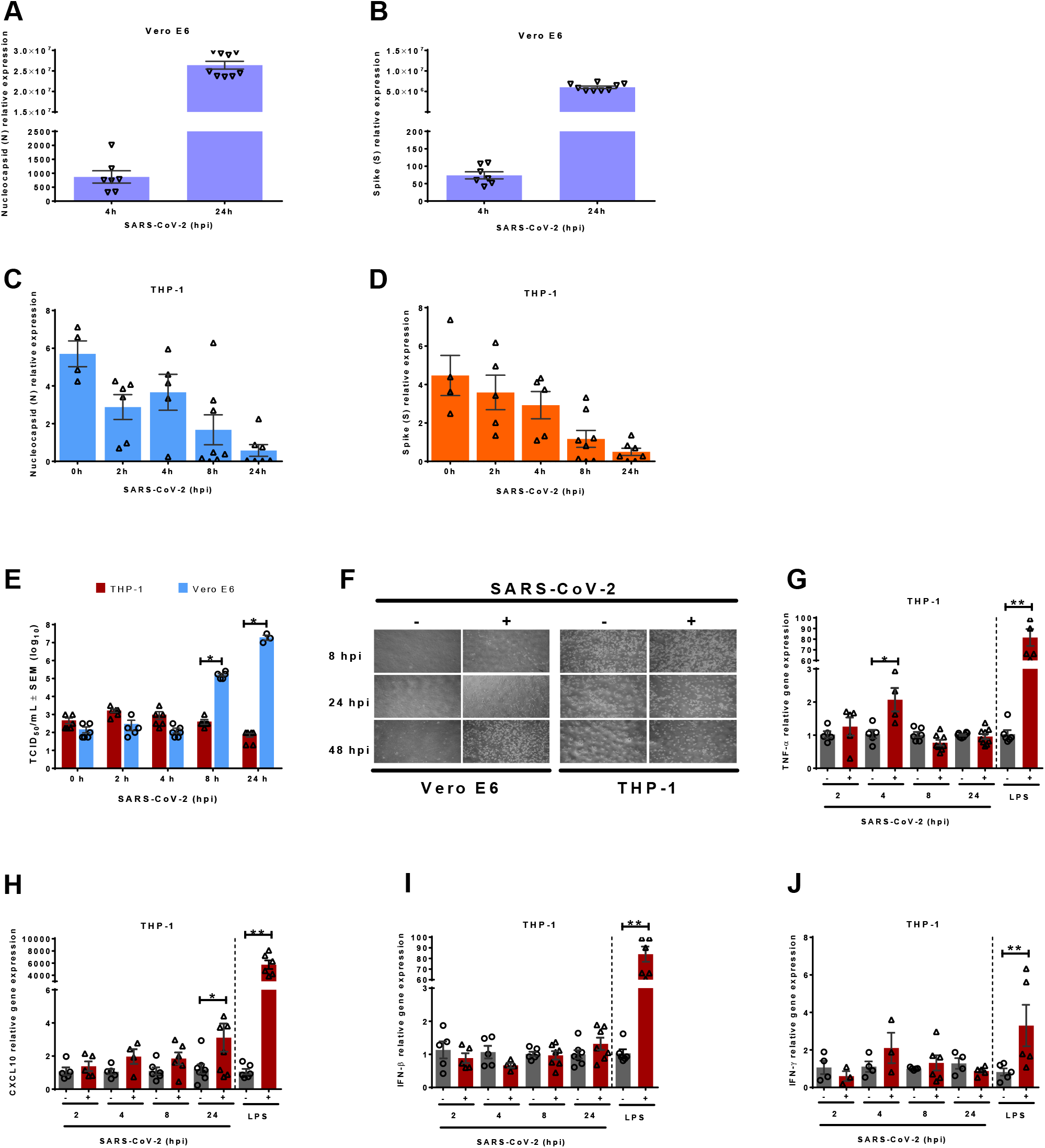
THP-1 macrophages express cytokines following SARS-CoV-2 infection in the absence of productive virus replication. RT-qPCR was used to detect SARS-CoV-2 N (A) and S (B) viral genes in mock infected or SARS-CoV-2 infected Vero E6 cells. RT-qPCR was used to detect N (C) and S (D) viral genes in mock infected or SARS-CoV-2 infected THP-1 macrophages. E) Culture supernatants from mock infected or SARS-CoV-2 infected THP-1 macrophages and Vero E6 cells were analyzed by TCID_50_ assay to determine infectious virus production. F) Bright field microscopy photographs of Vero E6 (MOI=0.1) and THP-1 (MOI=0.5) macrophages infected with SARS-CoV-2 for indicated times. RT-qPCR was used to detect relative gene expression of cytokines TNF-α (G), CXCL10 (H), IFN-β (I) and IFN-γ (J) in mock infected or SARS-CoV-2 infected THP-1 macrophages. LPS-treated macrophages (100ng/mL, 4 hours) were used as controls. Error bars denote the standard error of the mean (SEM) from 2 to 3 biologically independent experiments. Hpi= hours post infection. \**p*< 0.05 determined by Two-way ANOVA adjusted by Sidak’s multiple comparison test.

**Figure 3.**
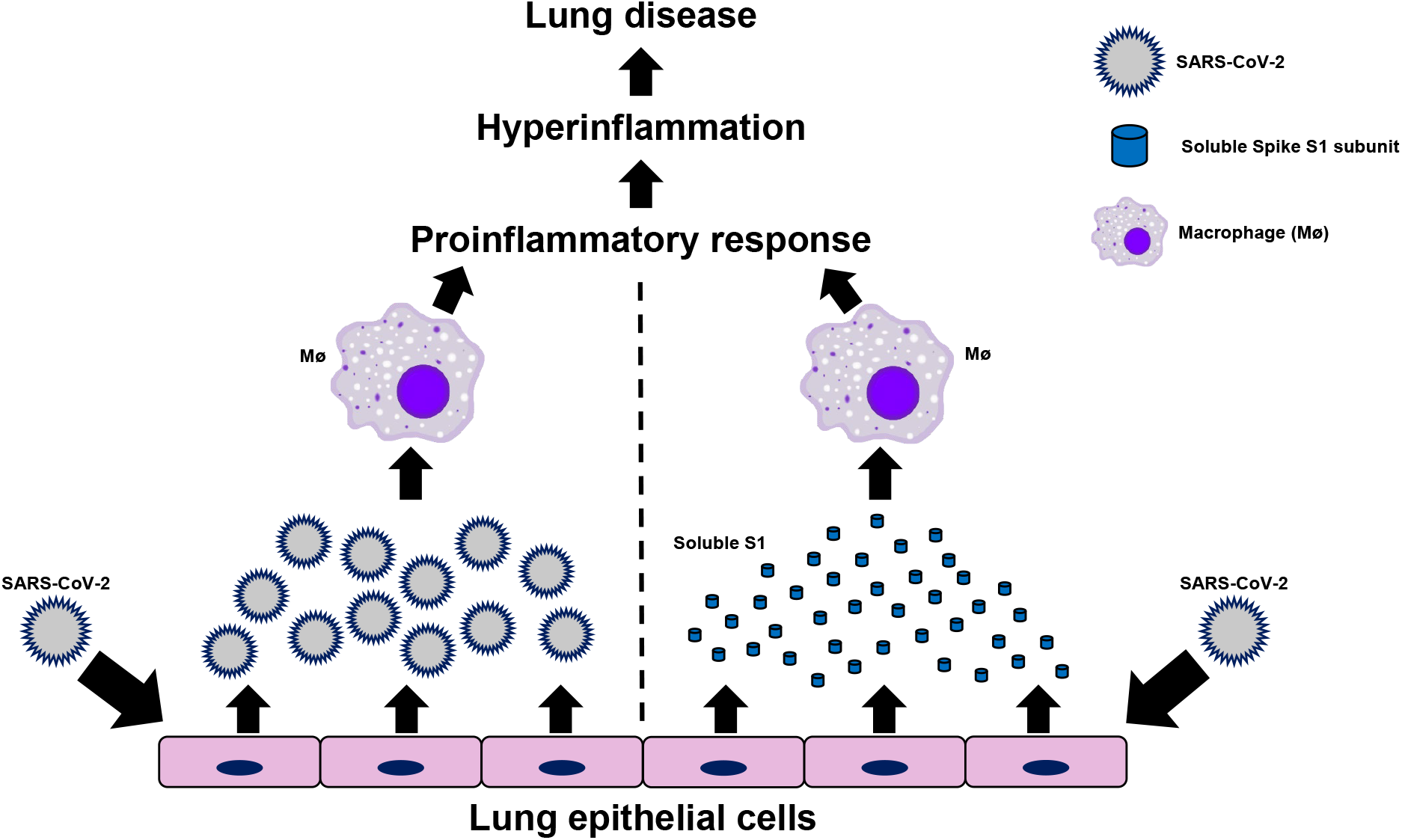
Proposed contribution of SARS-CoV-2 and soluble S1 in inducing inflammatory responses by macrophages. SARS-CoV-2 virions and soluble S1 proteins shed from productively infected epithelial cells trigger proinflammatory responses by macrophages, which then contribute to hyperinflammation and lung disease associated with COVID-19.

Despite non-productive replication, expression of TNF-α (Fig. 2G) and CXCL10 (Fig. 2H) in THP-1 macrophages infected with SARS-CoV-2 was significantly upregulated by 2- and 3-fold at 4 and 24 hpi, respectively. Additionally, SARS-CoV-2 infection did not induce IFN-β (Fig.2I) or IFN-γ (Fig. 2J) expression in THP-1 macrophages, whereas LPS upregulated both cytokines by 80- and 3-fold, respectively. Since both TNF-α and CXCL10 are key pro-inflammatory cytokines, our results suggest low-grade pro-inflammatory response in THP-1 macrophages infected with SARS-CoV-2 in the absence of productive virus replication. However, SARS-CoV-2 infection did not prompt type-I interferon dependent antiviral response in these cells.

Macrophages promote pro-inflammatory responses by producing cytokines and chemokines that initiate and sustain inflammation. The role of macrophages in SARS-CoV-2 infection remains unclear despite their function as pro-inflammatory cells and contribution to immune dysregulation. Our study shows that although THP-1 macrophages do not support productive virus replication, infection with SARS-CoV-2 upregulates expression of pro-inflammatory mediators linked to generalized hyperinflammation in COVID-19 patients^4,23^. Our study identifies the SARS-CoV-2 soluble S1 subunit as a viral factor involved in activation of pro-inflammatory response in macrophages. The soluble S1 subunit triggers pro-inflammatory response in non-infected macrophages and therefore, interaction with the S1 subunit suffices to induce this response independently of virus infection. Extracellular soluble S1 originating from infected lung epithelial cells may interact with uninfected macrophages to stimulate inflammation. Shedding of dissociated S1 has been shown during expression of the entire S protein on the surface of pseudoviruses^14^ and in mammalian cells^24^. Our study raises the possibility that cell-free soluble S1 engages with non-infected macrophages to induce low-grade pro-inflammatory response. SARS-CoV-2 infection in THP-1 macrophages does not lead to CPE (indicative of cell death) or productive viral replication, while low-grade inflammatory response remains upregulated for at least 24 hours after SARS-CoV-2 exposure (Fig. 2H). Therefore, virus-infected or S1-activated macrophages may become sources of pro-inflammatory mediators contributing to hyperinflammation in COVID-19 patients. Our study provides evidence for contribution of human macrophages to inflammation-associated immunopathology of SARS-CoV-2 by two possible mechanisms (graphical abstract) – a) virus infection of macrophages, and b) interaction of soluble S1 protein with uninfected macrophages.

## 3. Materials and Methods

### Cells and virus

Cell cultures were maintained at 37C in a 5% CO_2_ atmosphere. Vero E6 cells (ATCC; catalog no. CRL-1586) were cultured in DMEM medium (Gibco 12430062) supplemented with 10% FBS, 100 IU/ml Penicillin and 100 μg/ml Streptomycin. Human monocyte-like cells, THP-1 cell line (ATCC; catalog no. TIB-202), were cultured in RPMI 1640 medium (Gibco 21870076) supplemented with 10% FBS, 10mM HEPES, 1mM sodium pyruvate, 50μM of beta-mercaptoethanol, 100 IU/ml Penicillin and 100 μg/ml Streptomycin. SARS-CoV-2 isolate USA-WA1/2020 (BEI resources catalog no. NR-52281) was propagated in Vero E6 cells to generate a virus stock with a titer of 1.76 ×10^6^ 50% tissue culture infective doses (TCID_50_)/mL. All the SARS-CoV-2 titrations were performed by TCID_50_ assay on Vero E6 cells and titers were calculated by the method of Reed and Muench. Work with infectious virus was performed in biosafety cabinets within a biosafety containment level 3 facility. Personnel wore powered air purifying respirators during all procedures (MAXAIR Systems, Irvine, CA).

### Cell infection

THP-1 monocyte-like cells were seeded in the presence of phorbol 12-myristate 13-acetate (PMA, 100ng/mL) to induce differentiation into macrophages. After 24 hours of incubation, undifferentiated cells were washed away and remaining differentiated macrophages were incubated in fresh media without PMA for an additional 24 hours. THP-1 cells were either mock infected with supernatants from non-infected Vero E6 cells or inoculated at an MOI of 0.5 for 1 hour at 37C with virus stock generated in Vero E6 cells. Cells were washed once with PBS and incubated at 37°C in complete media for the indicated times. Vero E6 cells were seeded and incubated for 24 hours before virus infection at an MOI of 0.1 following the same procedure as with THP-1 cells. After indicated times, culture supernatants were collected for titration assays (TCID_50_) and RNA was extracted from infected cells using RNeasy Plus Mini Kit (Qiagen; catalog no. 74134) following manufacturer’s instructions. RNA extracted from THP-1 cells treated with 100ng/mL of LPS (Invivogen, catalog no. tlrl-eklps) for 4 hours was used as a control.

### Cell treatment

THP-1 monocyte-like cells were seeded in presence of phorbol 12-myristate 13-acetate (PMA, 100ng/mL) and allowed to differentiate as described above. Cells were treated with 8nM (0.6μg/mL) of recombinant soluble SARS-CoV-2 Spike S1 protein purified from HEK293 cells (SinoBiological; catalog no. 40591-V08H-B) or an equivalent volume of vehicle control for specified times. Cell culture supernatants were collected for ELISA assays and RNA was extracted from infected cells using Trizol (ThermoFisher; catalog no. 15596026) following manufacturer’s instructions.

### Reverse Transcription Quantitative PCR (RT-qPCR)

For cellular genes, total RNA (500ng) was used for cDNA synthesis using High-Capacity cDNA Reverse Transcription Kit (Applied Biosystems; catalog no. 4368814) following manufacturer’s instructions. Approximately 20ng of cDNA were used as template for qPCR reactions using SSOAdvanced Universal SYBR Green Supermix following manufacturer’s instructions (BioRad; catalog no.1725271). Following primers were used for detection of cellular genes by qPCR:

GAPDH Fw:5’-ACAACTTTGGTATCGTGGAAGG-3’; Rv: 5’-GCCATCACGCCACAGTTTC-3’
TNF-α Fw:5’-CCTCTCTCTAATCAGCCCTCTG-3’; Rv:5’- GAGGACCTGGGAGTAGATGAG-3’
IL6 Fw: 5’- ACTCACCTCTTCAGAACGAATTG-3’; Rv: 5’- CCATCTTTGGAAGGTTCAGGTTG-3’
IFN-γ Fw: 5’- TCGGTAACTGACTTGAATGTCCA-3’; Rv: 5’- TCGCTTCCCTGTTTTAGCTGC-3’
IFN-β Fw: 5’- GCTTGGATTCCTACAAAGAAGCA-3’; Rv: 5’-ATAGATGGTCAATGCGGCGTC-3’
CXCL10 Fw: 5’-GTGGCATTCAAGGAGTACCTC-3’; Rv: 5’- GCCTTCGATTCTGGATTCAGACA-3’

qPCR reactions were performed in a CFX96 Touch Real-Time PCR Detection System (Biorad, CA). Relative gene expression of target genes was determined using the average Ct for technical replicates normalized to GAPDH. Fold change over mock-infected cells was determined using 2^−ddCt^ method.

For quantification of viral RNA, total RNA (250ng or 500ng) was used for cDNA synthesis using SuperScript IV VILO Master Mix (Invitrogen, catalog no. 11756050) using the manufacturer’s instructions. cDNA was diluted to 1:10 and used as a template for qPCR reactions using PowerUp SYBR Green Master Mix (Applied Biosystems, catalog no. A25742). Primers for detection of SARS-CoV-2 genes^25^ N (Fw:5’-CAATGCTGCAATCGTGCTAC-3’; Rv:5’- GTTGCGACTACGTGATGAGG-3’) and S (Fw:5’- GCTGGTGCTGCAGCTTATTA-3’; Rv: 5’- AGGGTCAAGTGCACAGTCTA-3’) were used for qPCR, along with GAPDH primers listed above. qPCR reactions were performed using a StepOnePlus Real-Time PCR System (Applied Biosystems, CA), Ct values were determined using Design and Analysis 2.5.0 (Applied Biosystems), and normalized to GAPDH RNA levels using 2^−dCt^.

### Enzyme-linked immunosorbent assay (ELISA)

Human TNF alpha Uncoated ELISA Kit (ThermoFisher, catalog no. 88-7346) was used to determine secretion of TNF-α in culture supernatants of cells treated with SARS-CoV-2 S1 subunit. Cytokine concentration was calculated according to manufacturer’s instructions.

### Statistical analysis

Two-Way ANOVA adjusted by Sidak’s multiple comparison test was performed to evaluate relative expression from RT-qPCR data and ELISA from three experimental groups compared at multiple time points. A *p* value of <0.05 was considered significant for all statistical tests. All statistical tests were performed using GraphPad Prism v6.01 (CA, USA).

## Authorship

T.A.M., K.C., K.H. and L.G.M. performed the experiments. T.A.M., K.C. and S.B. contributed to the experimental design, data analyses and interpretation, preparation of figures and tables, and prepared the manuscript. All authors have approved the final manuscript as submitted and agree to be accountable for all aspects of the study.

## Acknowledgements

This research was supported by funding from the Washington Research Foundation (SB) and National Institutes of Health (NIH) R01AI083387 (SB). LGM was supported by NIH NIGMS Predoctoral Biotechnology Training Grant 5T32GM008336. The authors thank Sedelia Dominguez and Shannon Allen at Washington State University’s Paul G. Allen School for Global Health for their assistance in the BSL3 facility.

